# TAEL 2.0: An Improved Optogenetic Expression System for Zebrafish

**DOI:** 10.1101/2020.09.24.312470

**Authors:** Jesselynn LaBelle, Adela Ramos-Martinez, Kyle Shen, Laura B. Motta-Mena, Kevin H. Gardner, Stefan C. Materna, Stephanie Woo

**Affiliations:** Department of Molecular Cell Biology, University of California, Merced, CA 95343 USA; Optologix, Inc., Dallas, TX 75201 USA; Structural Biology Initiative, Advanced Science Research Center, CUNY; New York, NY, 10031, USA; Department of Chemistry & Biochemistry, City College of New York, New York, USA; Ph.D. Programs in Biochemistry, Biology, and Chemistry, The Graduate Center, City University of New York, New York, USA

**Keywords:** Zebrafish, Optogenetics, Gene expression, Light-activated transcription factor

## Abstract

Inducible gene expression systems are valuable tools for studying biological processes. We previously developed an optogenetic gene expression system called TAEL that is optimized for use in zebrafish. When illuminated with blue light, TAEL transcription factors dimerize and activate gene expression downstream of the TAEL-responsive C120 promoter. By using light as the inducing agent, the TAEL/C120 system overcomes limitations of traditional inducible expression systems by enabling fine spatial and temporal regulation of gene expression. Here, we describe ongoing efforts to improve the TAEL/C120 system. We made modifications to both the TAEL transcriptional activator and the C120 regulatory element, collectively referred to as TAEL 2.0. We demonstrate that TAEL 2.0 consistently induces higher levels of reporter gene expression and at a faster rate, but with comparable background and toxicity as the original TAEL system. With these improvements, we were able to create functional stable transgenic lines to express the TAEL 2.0 transcription factor either ubiquitously or with a tissue-specific promoter. We demonstrate that the ubiquitous line in particular can be used to induce expression at late embryonic and larval stages, addressing a major deficiency of the original TAEL system. This improved optogenetic expression system will be a broadly useful resource for the zebrafish community.

## Introduction

Inducible gene expression systems are valuable tools for studying biological processes as they enable user-defined control over the timing, location, and level of expression. In zebrafish and other model organisms, the most widely used inducible expression systems fall into two broad categories – those that rely on the heat shock response^1^ and those using small molecule inducing agents^2^. More recently, optogenetic approaches have been developed based on light-sensitive transcription factors^3–6^. One such system is based on EL222, a naturally occurring blue light-activated transcription factor found in the bacterium *Erythrobacter litoralis HTCC2594*. The endogenous transcription factor contains a light-oxygen-voltage-sensing (LOV) domain that in response to blue light (450 nm) undergoes a conformational change and dimerizes, allowing it to bind and initiate transcription from a regulatory element termed C120^7^. EL222 was the basis for an inducible expression system designed for mammalian cell culture^8^. Our group previously designed EL222 for use in zebrafish by fusing it to a KalTA4 transcriptional activation domain, which minimized toxicity in zebrafish embryos while still maintaining functionality^6^. We demonstrated that this KalTA4-EL222 fusion protein, which we termed TAEL, could be combined with C120-containing transgenes to achieve light-inducible expression of multiple genes of interest. We also validated multiple approaches for delivering patterned blue light illumination to spatially and temporally control induction in zebrafish embryos. However, we were unable to establish stable transgenic lines for TAEL expression that could induce expression from our C120 reporter lines, suggesting that TAEL and/or the C120 promoter could be further optimized.

In this study, we present ongoing efforts to improve the function of the TAEL/C120 system. We made changes to both the TAEL transcriptional activator and the C120 promoter, collectively termed TAEL 2.0, that produce significantly higher levels of light-induced expression at a faster rate. Importantly, these improvements allowed us to address a major deficiency of our previously published system (referred to here as TAEL 1.0), namely the lack of functional, stable transgenic lines for both TAEL and C120 components. Here, we describe the generation of transgenic lines that express functional TAEL 2.0 components either ubiquitously or in the developing endoderm. We demonstrate that the ubiquitous line in particular can be used to induce expression at late embryonic and larval stages, extending the use of this system beyond early embryo stages.

## Materials and Methods

### Vector construction and mRNA synthesis

#### pμTol2 backbone

For expression plasmids and transgenes created for this study, we generated a minimal plasmid backbone called pμTol2, which can be used for both Tol2-based transgenesis and in vitro mRNA synthesis. Its short length of 2520 base pairs enables modification of inserts by PCR through the backbone, thus eliminating the need to subclone. In brief, pμTol2 was constructed by Gibson assembly, fusing the Tol2 sites for genomic integration^9^ with the commonly used expression cassette of pCS2 including polylinkers and SV40 polyadenylation site^10,11^ and a plasmid backbone derived from pUC19^12^. To ensure efficient protein synthesis, all plasmids newly constructed for this study contain the zebrafish-optimized Kozak sequence 5’-GCAAACatgG-3’, where the lower case “atg” denotes the start codon^13^.

#### Expression plasmids

pCS2-TAEL has been described previously^6^. To construct expression plasmids pμTol2-N-TAEL, Optologix, Inc. (Dallas, TX) provided synthesized oligomers containing the SV40 large T-antigen nuclear localization signal. We fused these to the 5’ end of the TAEL ORF and to the pμTol2 backbone by Gibson assembly^14^. Similarly, pμTol2-TAEL-N was constructed by fusing synthesized oligomers containing the nucleoplasmin nuclear localization signal (also provided by Optologix, Inc.) to the 3’ end of the TAEL ORF by Gibson assembly^14^. Capped messenger RNA was synthesized using mMESSAGE mMACHINE SP6 kit (Ambion) with plasmids cut with NotI as linear template. For experiments in Fig. 1–3, Tg(C120:mCherry;cryaa:Venus) or Tg(C120F:mCherry) males were crossed to wild-type females and resulting embryos were each injected with ~50 pg of TAEL, N-TAEL, or TAEL-N mRNA at the 1-cell stage.

**Figure 1.**
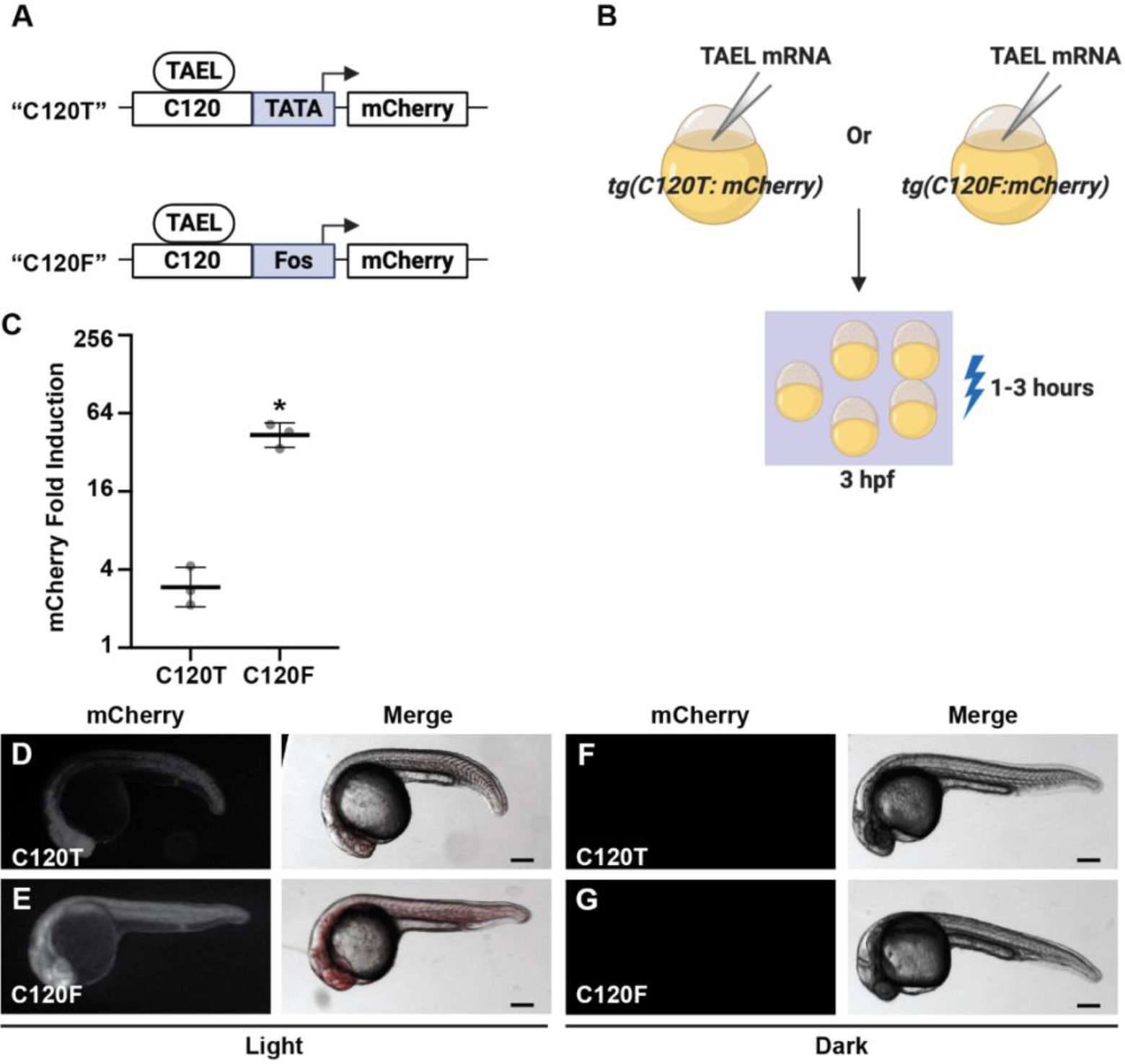
Coupling the C120 regulatory element to the Fos basal promoter significantly increases light-induced expression. **A.** Schematic comparing different C120-based reporter constructs in which TAEL-responsive C120 sequences (C120) were coupled to either a minimal TATA box (TATA) or the basal promoter from the mouse *Fos* gene (Fos) and used to drive expression of mCherry. **B.** Schematic of experimental design. *Tg(C120T:mCherry)* or *Tg(C120F:mCherry)* embryos were injected with TAEL mRNA. mCherry expression was induced by illuminating embryos with blue light starting at 3 hours post-fertilization (hpf). **C.** Comparison of light-induced mCherry expression in *Tg(C120T:mCherry)* and *Tg(C120F:mCherry)* embryos injected with TAEL mRNA. mCherry transcript levels were measured by qPCR from embryos illuminated with blue light for 1 hour and compared to sibling embryos kept in the dark. Y-axis is set at log_2_ scale. Dots represent biological replicates. Solid lines represent mean. Error bars represent S.D. *p<0.05. **D-G.** Representative images of mCherry fluorescence in *Tg(C120T:mCherry)* (D, F) or *Tg(C120F:mCherry)* (E, G) embryos injected with TAEL mRNA and illuminated with blue light for 3 hours (D, E) or kept in the dark (F, G). Images were acquired between 20 and 24 hours post-illumination. Scale bars, 200 μm.

**Figure 3.**
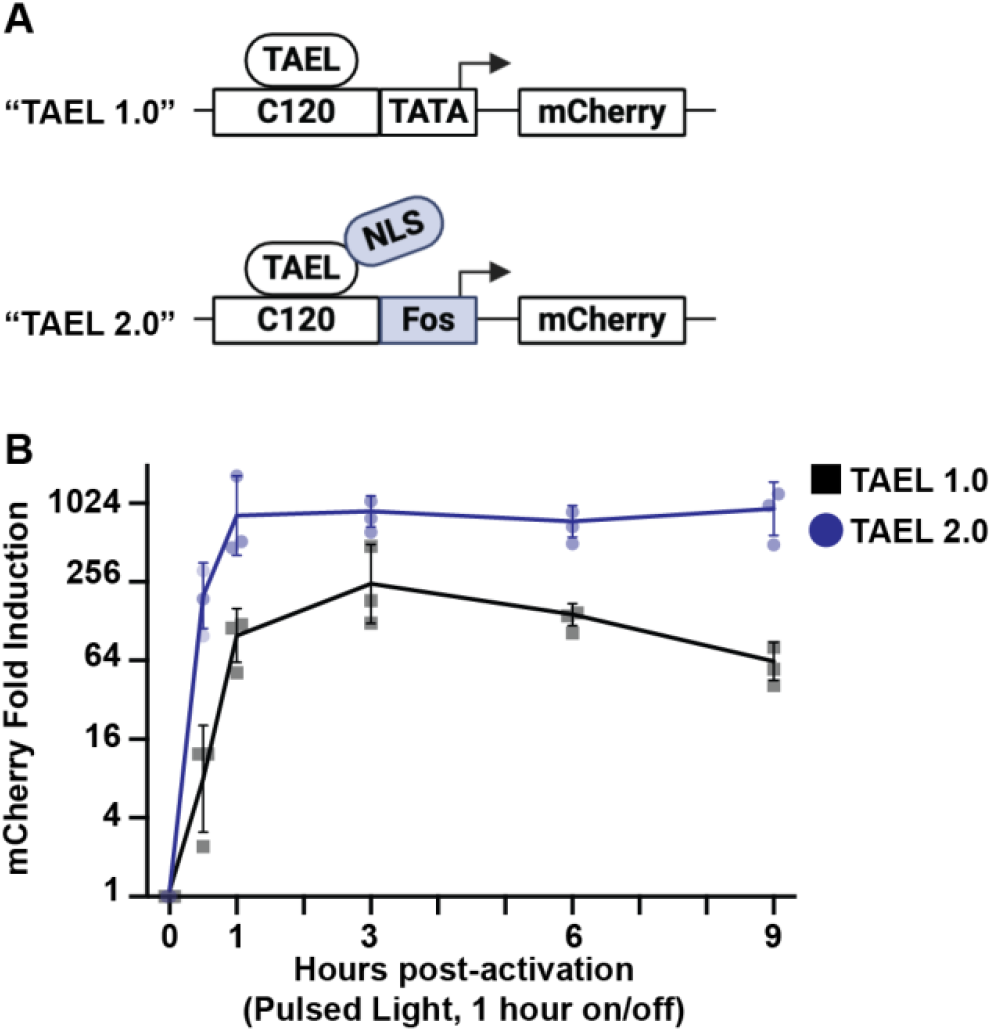
TAEL 2.0 modifications improve both the rate and level of light-induced expression. **A**. Schematic comparing TAEL 1.0 and TAEL 2.0. TAEL 1.0 consists of the TAEL transcription factor that lacks an NLS and the C120T promoter containing a minimal TATA box sequence. TAEL 2.0 consists of the TAEL-N transcription factor with a C-terminal NLS and the C120F promoter containing the basal promoter from the mouse *Fos* gene. **B.** Comparison of light-induced mCherry expression over time using TAEL 1.0 (black, dots) or TAEL 2.0 (blue, squares). *Tg(C120T:mCherry)* or *Tg(C120F:mCherry)* embryos were injected with mRNA for TAEL or TAEL-N, respectively. mCherry expression was activated by illuminating embryos with blue light (pulsed at a frequency of 1 hour on/1 hour off), starting at 3 hours post-fertilization. mCherry transcript levels were measured by qPCR at the indicated time points and normalized to 0 h post-activation. Y-axis is set at log_2_ scale. Dots and squares represent biological replicates. Solid lines represent mean. Error bars represent S.D.

#### Transgene plasmids

To construct pμTol2-C120F:mCherry, the mouse *Fos* basal promoter sequence: 5’-CCAGTGACGTAGGAAGTCCATCCATTCACAGCGCTTCTATAAAGGCGCCAGCTGAGGCGCCTACTACTCCAACCGCGACTGCAGCGAGCAACT-3’^15^ was synthesized by Integrated DNA Technologies and the C120 sequence^6^ was amplified by PCR. These sequences were fused together and inserted into pμTol2 by Gibson assembly. The transgene plasmid pμTol2-C120F:GFP was constructed by separate PCR amplification of the C120F promoter and GFP ORF which were then cloned into pμTol2 by Gibson assembly. pμTol2-sox17:TAEL-N was constructed by separate PCR amplification of the *sox17* promoter^16^ and TAEL-N ORF which were then cloned into pμTol2 by Gibson assembly. pμTol2-ubb:TAEL-N was constructed by separate PCR amplification of the *ubb* promoter^17^ and TAEL-N ORF, which were then cloned into pμTol2 by Gibson assembly.

All plasmids constructed for this study are available by direct request.

### Zebrafish Strains

Adult Danio rerio zebrafish were maintained under standard laboratory conditions. Zebrafish in an outbred AB, TL, or EKW background were used as wildtype strains. *Tg(C120:mCherry;cryaa:Venus)*^*sfc14*^, referred to here as *Tg(C120T:mCherry)*, has been previously described^6^. *Tg(C120-Mmu.Fos:mCherry)*^*ucm104*^, *Tg(C120-Mmu.Fos:GFP)^ucm107^*, *Tg(ubb:TAEL-N)*^*ucm113*^, and *Tg(sox17:TAEL-N)*^*ucm114*^ were generated using standard transgenesis protocols^9,18^. This study was performed with the approval of the Institutional Animal Care and Use Committee (IACUC) of the University of California Merced.

### Global light induction

Global light induction was provided by a MARS AQUA-165-55 110W LED aquarium hood. Actual power of light received by embryos (lids of plates removed) was measured as ∼1.6 mW/ cm^2^ at 456 nm. For experiments in Fig. 1–2, 4 hpf (hours post-fertilization) embryos were illuminated with constant blue light for 1–3 hours. For experiments in Fig. 3–6, a timer was used to apply constant or pulsed light (NEARPOW Timer Switch). Dark controls were placed in a light proof box in the same 28.5°C incubator as the light-treated samples.

**Figure 2.**
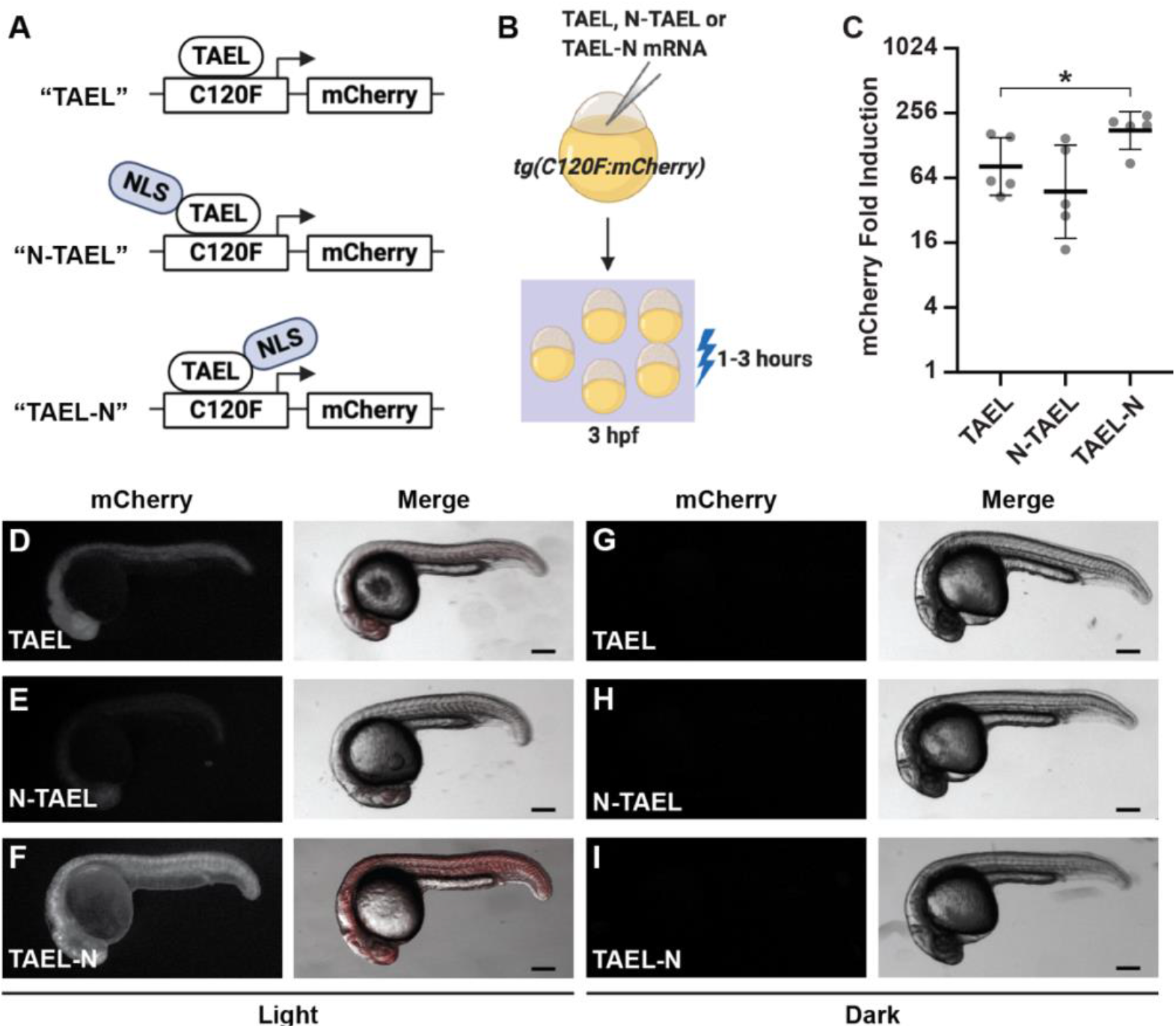
Adding a C-terminal nuclear localization signal (NLS) to TAEL significantly increases light-induced expression. **A.** Schematic comparing different TAEL constructs containing no NLS (TAEL), one N-terminal NLS (N-TAEL), or one C-terminal NLS (TAEL-N). **B.** Schematic of experimental design. *Tg(C120F:mCherry)* embryos were injected TAEL, N-TAEL, or TAEL-N mRNA. mCherry expression was induced by illuminating embryos with blue light starting at 3 hours post-fertilization (hpf). **C.** Comparison of light-induced mCherry expression in *Tg(C120F:mCherry)* embryos injected with TAEL, N-TAEL, or TAEL-N mRNA. mCherry transcript levels were measured by qPCR from embryos illuminated with blue light for 1 hour and compared to sibling embryos kept in the dark. Y-axis is set at log_2_ scale. Dots represent biological replicates. Solid lines represent mean. Error bars represent S.D. *p<0.05. **D-I.** Representative images of mCherry fluorescence in *Tg(C120F:mCherry)* embryos injected with TAEL (D, G), N-TAEL (E, H), or TAEL-N (F, I) mRNA and illuminated with blue light for 3 hours (D-F) or kept in the dark (G-I). Images were acquired between 20 and 24 hours post-illumination. Scale bars, 200 μm.

**Figure 4.**
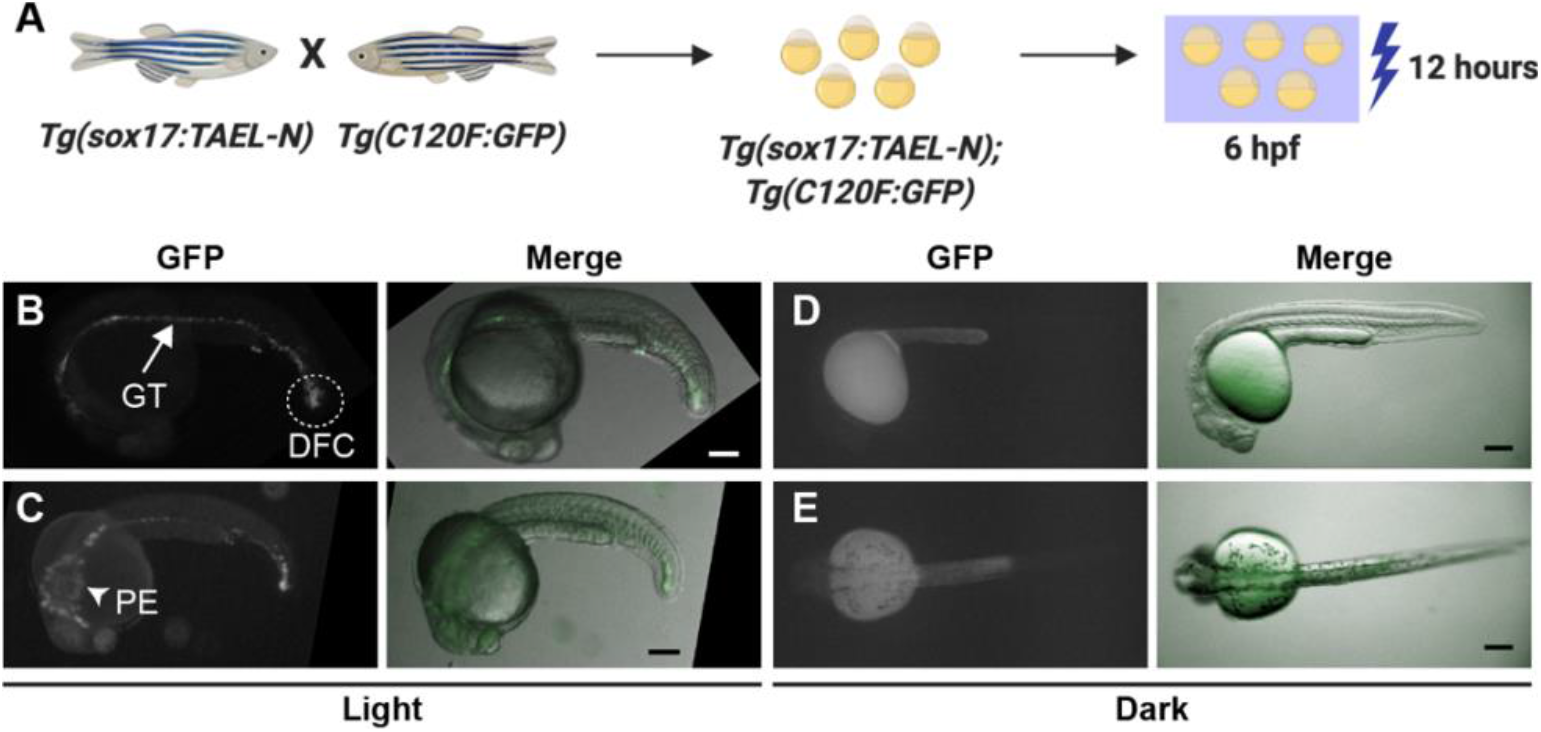
The stable transgenic line *Tg(sox17:TAEL-N)* restricts light-induced expression to endoderm-derived tissues. **A** Schematic depicting experimental design. *Tg(sox17:TAEL-N)* and *Tg(C120F:GFP)* adult zebrafish were crossed to produce double transgenic embryos. GFP expression was activated by illuminating embryos for 12 hours, starting at 6 hours post-fertilization (hpf), with blue light pulsed at a frequency of 1 hour on/1 hour off. **B-E.** Representative images of *Tg(sox17:TAEL-N);Tg(C120F:GFP)* embryos exposed to blue light (B-C) or kept in the dark (D-E). Images were acquired between 18 and 20 hours post-activation. Arrow in (B) indicates gut tube (GT). Dashed lines in (B) indicate derivatives of the dorsal forerunner cells (DFC). Arrowhead in (C) indicates pharyngeal endoderm (PE). B, D are lateral views, anterior to the left. C, E are dorsal views, anterior to the left. Scale bars, 200 μm.

**Figure 6.**
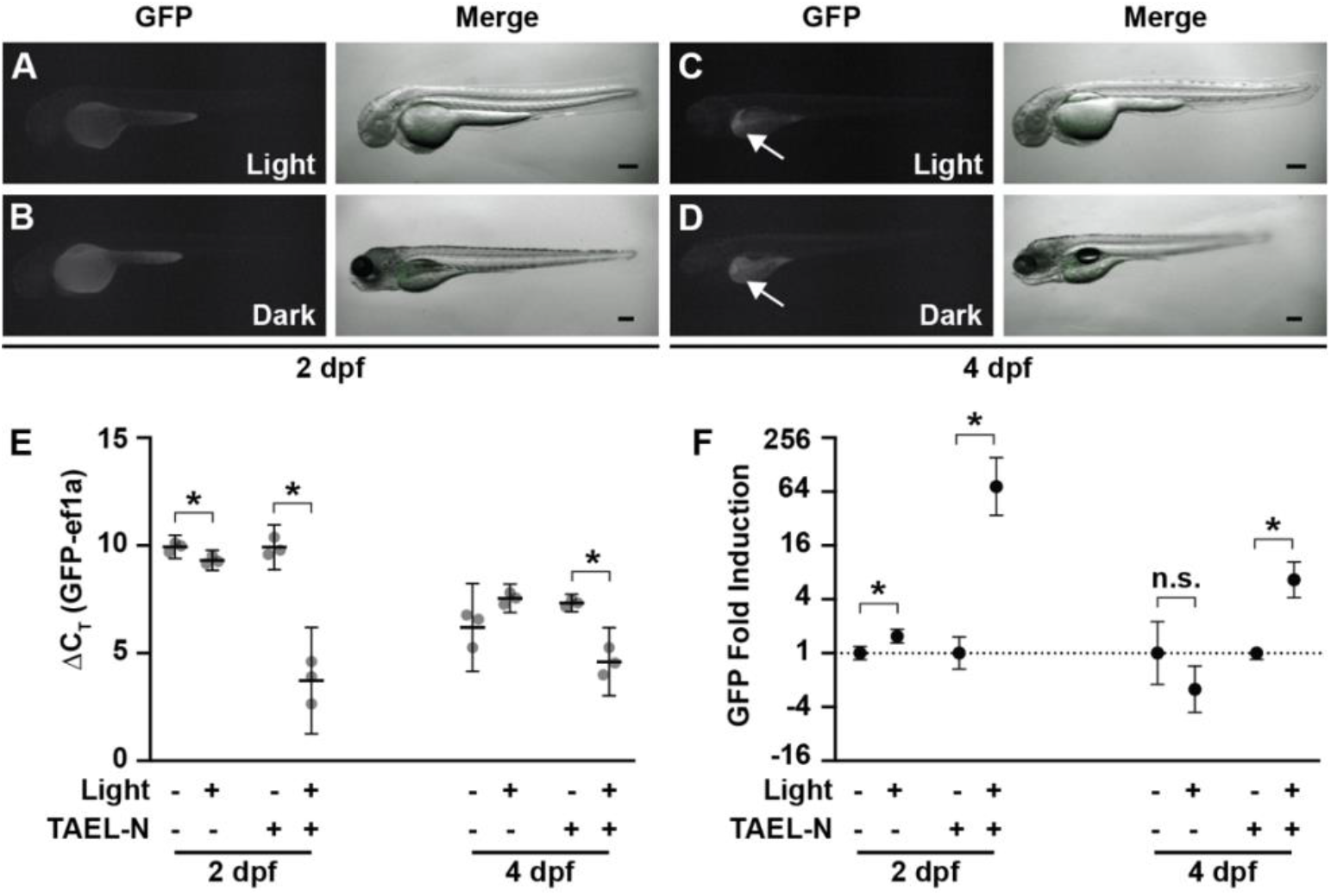
Basal expression from *Tg(C120F:GFP)* is not responsive to light. **A-D.** Representative images of *Tg(C120:GFP)* embryos at 2 days post-fertilization (dpf) (A-B) or larvae at 4 dpf (C-D). Embryos were illuminated for 12 hours with blue light pulsed at a frequency of 1 hour on/1 hour off (A, C) or kept in the dark (B, D). Images were acquired between 18 and 20 hours after exposure to the pulsed light regime. Arrows in (C, D) point to ectopic liver expression of GFP. Scale bars, 200 nm. **E.** qPCR analysis of GFP expression from *Tg(C120:GFP)* or *Tg(C120F:GFP);Tg(ubb:TAEL-N)* zebrafish at 2 or 4 dpf illuminated with constant blue light for 1 hour or kept in the dark. Data are presented as ΔC_T_ values normalized to the housekeeping gene *ef1a*. Dots represent biological replicates. Solid lines represent mean. Error bars represent S.D. *p<0.05. **F.** Fold induction of GFP expression in response to light calculated from the same qPCR analysis shown in (E). Y-axis is set at log_2_ scale. Data are presented as mean ± S.D. *p<0.05. n.s., not significant.

### Real-time quantitative PCR

To quantify light-induced expression, total RNA from 30–50 light-treated or dark control embryos was extracted using the illustra™ RNAspin Mini kit (GE Healthcare). 1 μg total RNA was used for reverse transcription with qScript XLT cDNA SuperMix (Quantabio). Each qPCR reaction contained 2X PerfeCTa^®^ SYBR green fast mix (Quantabio), 5-fold diluted cDNA and 325 nM each primer. Reactions were carried out on a QuantStudio3 (Applied Biosystems) real time PCR machine using the following program: initial activation at 95°C for 10 min, followed by 40 cycles of 30 s at 95°C, 30 s at 60°C and 1 min at 72°C. Once the PCR was completed, a melt curve analysis was performed to determine reaction specificity. Data represent averages from 3–5 biological replicates, each with three technical replicates. The housekeeping gene *ef1a* was used as a reference. Fold change was calculated using the 2^(−ΔΔCT)^ method^19^. Statistical significance was determined by Welch’s t-test (unless otherwise stated) using Prism software (GraphPad). qPCR primers used are: mcherry forward: 5’-GACCACCTACAAGGCCAAGA-3’; mcherry reverse: 5’-CTCGTTGTGGGAGGTGATGA-3’; ef1a forward 5’-CACGGTGACAACATGCTGGAG-3’; ef1a reverse: 5’-CAAGAAGAGTAGTACCGCTAGCAT-3’

### Microscopy and image processing

Fluorescence and brightfield images were acquired on a Leica dissecting stereomicroscope or Olympus dissecting stereomicroscope. Dechorionated embryos or larvae were embedded in 1.5% low-melting agarose (ISC BioExpress) containing 0.01% tricaine (Sigma-Aldrich) within glass-bottom Petri dishes (MatTek Corporation). Standard filter settings were applied and brightfield and fluorescence images were merged after acquisition. Identical exposure settings for fluorescence images were used for all embryos from the same set of experiments. All image processing was performed using ImageJ software^20^. Illustrations were created with BioRender (https://biorender.com/).

## Results

### TAEL-induced expression is increased by coupling the C120 regulatory element to a *Fos* basal promoter

In our previously published system, the TAEL-responsive C120 regulatory sequence was coupled to a minimal TATA box^6,8^. Because this minimal TATA box originated from a mammalian expression vector, we reasoned that using a zebrafish-optimized basal promoter instead would improve performance of the TAEL system. The basal promoter from the mouse *Fos* gene was previously shown to function well in zebrafish transgenes, allowing for high expression levels with minimal background^15,21^. Therefore, we constructed a new TAEL-responsive promoter consisting of 5 repeats of the C120 regulatory sequence coupled to the mouse *Fos* basal promoter (*C120-Mmu.Fos*, abbreviated throughout as *C120F*). We then determined whether this new C120 promoter improves light-induced expression compared to the previous TATA box-containing version (Fig. 1). First, we generated a stable transgenic zebrafish line using *C120F* to control expression of an mCherry reporter (*Tg(C120F:mCherry)*) to make direct comparisons to our previously published reporter line^6^, referred to here as *Tg(C120T:mCherry)*. We injected both *Tg(C120T:mCherry)* and *Tg(C120F:mCherry)* embryos with ~50 pg TAEL mRNA then globally illuminated them with blue light starting at 3 hpf. qPCR analysis showed that compared to sibling control embryos kept in the dark, mCherry expression was induced 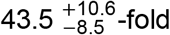 in *Tg(C120F:mcherry)* embryos, which was significantly higher than the 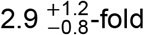 induction in *Tg(C120T:mCherry)* embryos (p=0.0009) (Fig. 1C). Consistent with these results, mCherry fluorescence was qualitatively brighter in *Tg(C120F:mCherry)* embryos compared to *Tg(C120T:mCherry)* embryos (Fig. 1D-E). Importantly, we did not observe mCherry fluorescence in embryos kept in the dark for either genotype (Fig. 1F-G). Together, these results suggest that coupling the C120 regulatory element with a *Fos* basal promoter instead of a minimal TATA box significantly increases TAEL-induced gene expression while maintaining low background expression.

### TAEL-induced expression is increased by adding a C-terminal nuclear localization signal to TAEL

Our original TAEL construct consists of a Kal-TA4 transcription activation domain, the light-sensitive LOV domain, and a DNA-binding domain that recognizes the C120 sequence but does not contain an explicit nuclear localization signal (NLS). Although TAEL can likely enter the nucleus through diffusion, because of its relatively small size of 257 amino acids, we wanted to test whether targeting TAEL specifically to the nucleus by adding an NLS would increase the amplitude of induction and improve light-induced expression (Fig. 2).

We first generated a construct in which the SV40 large T-antigen NLS was fused to the amino terminus of TAEL (N-TAEL). When delivered by mRNA injection into *Tg(C120F:mCherry)* embryos, we were surprised to find that N-TAEL induced mCherry expression less strongly (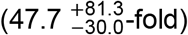) than the original TAEL protein (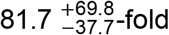; p=0.3398) (Fig. 2C). Consistent with these qPCR results, mCherry fluorescence was more variable and often dimmer in embryos injected with N-TAEL versus TAEL mRNA (Fig. 2D-E). We speculated that fusing the NLS to the N-terminus of TAEL places it directly adjacent to the KalTA4 transcriptional activation domain, which may negatively interfere with transactivation. Therefore, we generated a construct in which the nucleoplasmin NLS was fused to the carboxy terminus of TAEL (TAEL-N). By qPCR analysis, *Tg(C120F:mCherry)* embryos injected with TAEL-N mRNA showed higher levels of mCherry induction (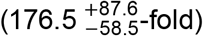) compared to both TAEL (p=0.053) and N-TAEL (p=0.0392) (Fig. 2C). As expected, mCherry fluorescence was brightest in embryos injected with TAEL-N (Fig. 2F). We did not observe mCherry fluorescence in any injected embryos kept in the dark (Fig. 2G-I). Together, these results demonstrate that adding a nuclear localization signal at the C-terminus of TAEL further increases light-induced gene expression with minimal background.

### TAEL 2.0 induces higher expression levels at a faster rate

We next characterized the effects of combining the modifications we made to the C120 promoter and TAEL transcriptional activator. With our previously published TAEL system, we found that sustained activation of gene expression over several hours could be achieved by pulsing the activating blue light at 1 hour on/off intervals^6^; under this illumination regime, peak expression levels were reached by 3 hours post-activation and sustained for up to 8 hours. To determine if TAEL 2.0 improves the kinetics and/or range of light-induced expression, we performed a similar time course of mCherry induction comparing *Tg(C120T:mCherry)* embryos injected with TAEL mRNA (“TAEL 1.0”) to *Tg(C120F:mcherry)* embryos injected with TAEL-N mRNA (“TAEL 2.0”). Starting at approximately 3 hpf, mCherry expression was activated by globally illuminating embryos with pulsed blue light (1 hour on, 1 hour off), and mCherry expression levels were measured by qRT-PCR at various timepoints up to 9 hours post-activation (Fig. 3). Throughout the entire time course, we found that TAEL 2.0 induced significantly higher mCherry expression compared to TAEL 1.0 (2-way ANOVA, p<0.0001). We further found that the rate of induction also improved. At 1 hour post-activation, mCherry expression had increased by 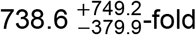 with TAEL 2.0 compared to 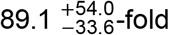 with TAEL 1.0. Finally, we found that these high levels of induction with TAEL 2.0 could be sustained for up to 9 hours under pulsed (1 hour on/off) illumination. Together, these results demonstrate that the combined modifications we made to the TAEL system improve both the range and induction kinetics of this light-activated expression system.

### TAEL 2.0 modifications enable functional stable transgenic lines of TAEL components

One notable deficiency of our previous TAEL system was the lack of functional stable transgenic lines expressing the TAEL transcriptional activator. With its greatly increased amplitude and kinetics of induction, we determined whether TAEL 2.0 could address this previous limitation.

We generated a stable transgenic line, *Tg(sox17:TAEL-N)*, to express TAEL-N under the *sox17* promoter, which drives expression in the endoderm and dorsal forerunner cells (DFCs)^16^. We crossed this line with a *Tg(C120F:GFP)* reporter line. The resulting double transgenic embryos were globally illuminated with pulsed blue light (1 hour on/off) or kept in the dark from 6–18 hpf (Fig. 4A). We observed GFP fluorescence in derivatives of the endoderm such as the gut tube and the pharyngeal endoderm as well as derivatives of the dorsal forerunner cells (DFCs) within the tail mesoderm in illuminated embryos but not those kept in the dark (Fig. 4B-E). Because activating blue light was applied globally, this result suggested that TAEL-N functions in, and is restricted to, the *sox17* expression domain. Additionally, we observed that the intensity of GFP fluorescence was brightest in the tail (Fig. 4B-C), again consistent with the known *sox17* expression pattern, which is highest in the DFCs. In sum, we successfully generated a stable transgenic line to express TAEL-N that enables tissue-specific induction of a gene of interest even when activating blue light is applied globally.

One consequence of the lack of functional stable transgenic lines for TAEL 1.0 is that its use is limited to early embryonic stages. To determine if TAEL 2.0 modifications could expand the range of accessible developmental stages, we generated a stable transgenic line, *Tg(ubb:TAEL-N)*, to express TAEL-N under the *ubb* promoter. This promoter has been shown to drive ubiquitous expression at all developmental stages^17^. We crossed this line to *Tg(C120F:GFP)* then exposed double transgenic embryos to activating blue light at several different time points spanning embryonic to larval stages (Fig. 5A). In all cases, we observed increased GFP fluorescence in illuminated embryos or larva but not in control siblings that had been kept in the dark (Fig. 5B-G).

**Figure 5.**
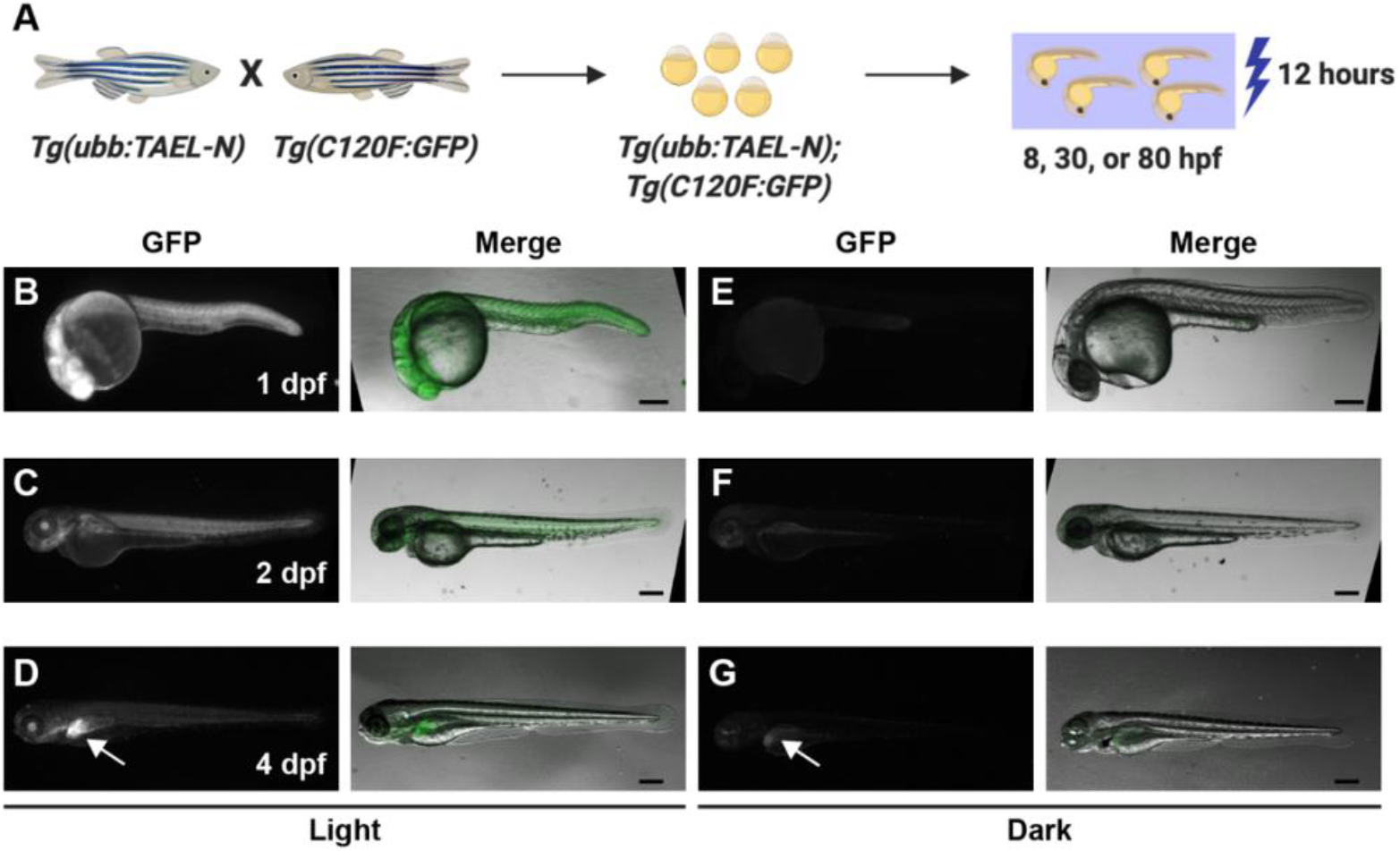
The stable transgenic line *Tg(ubb:TAEL-N)* enables light-induced expression at multiple developmental stages. A. Schematic depicting experimental design. *Tg(ubb:TAEL-N) and Tg(C120F:GFP)* adult zebrafish were crossed together to produce double transgenic embryos. GFP expression was activated at multiple time points by illuminating embryos for 12 hours with blue light pulsed at a frequency of 1 hour on/1 hour off. hpf, hours post-fertilization. **B-G.** Representative images of *Tg(ubb:TAEL-N);Tg(C120F:GFP)* embryos or larvae exposed to blue light (B-D) or kept in the dark (E-G). Images were acquired at the indicated stages between 18 and 20 hours post-activation. dpf, days post-fertilization. Arrows in (D, G) point to ectopic liver expression of GFP. Scale bars, 200 nm.

At 4 days post-fertilization (dpf), we observed GFP fluorescence in the livers of both illuminated and control larvae (arrows, Fig. 5D, G). This pattern appears specific to this *Tg(C120F:GFP)* line (when generating the line, we were only able to recover one founder with germline transmission of the transgene), and thus it is likely due to positional effects of transgene insertion. Our other zebrafish lines, including *Tg(C120F:mCherry* (data not shown), do not display similar fluorescence indicating it is unlikely due to autofluorescence. We could still visually detect light-dependent GFP induction above this background expression (Fig. 5D), suggesting that despite its ectopic liver expression, the *C120F:GFP* reporter transgene is functional at 4 dpf. Taken together, these results demonstrate that TAEL 2.0 can be used to induce expression in a broad range of developmental stages.

### Fidelity of the TAEL 2.0 system

A recent study showed that blue light alone can increase expression of *Fos* and other activity-dependent genes in cultured mouse cortical neurons^22^. Because the *C120F* promoter utilizes the basal promoter from the mouse *Fos* gene, it is possible that there are endogenous factors, especially in neural tissues, that can drive light-responsive expression from the *C120F* promoter. Such induction, independent of TAEL-N, would reduce the specificity of the TAEL 2.0 system.

To determine whether the *C120F* promoter is activated in the absence of TAEL-N, we exposed *Tg(C120F:GFP)* zebrafish to blue light at 2 dpf or 4 dpf; the latter time point was chosen as light-driven neuronal activity likely increases over time. Apart from the ectopic liver expression at 4 dpf described above, we did not observe appreciable GFP fluorescence in any animals (Fig. 6A-D).

To better uncover any TAEL-N-independent function of the *C120F* promoter, we used qPCR to quantify and compare GFP expression in *Tg(C120F:GFP)* and *Tg(C120F:GFP);Tg(ubb:TAEL-N)* animals. At 2 dpf, we detected equally low levels of GFP expression in both *Tg(C120F:GFP)* and *Tg(C120F:GFP);Tg(ubb:TAEL-N)* embryos kept in the dark. The average GFP ΔC_T_ values, normalized to the housekeeping gene *ef1a*, were 9.93±0.22 for *Tg(C120F:GFP)* and 9.91±0.42 for *Tg(C120F:GFP);Tg(ubb:TAEL-N)* (Fig. 6E). Although these low levels are insufficient to produce visible amounts of GFP (see Fig. 6B), they suggest that the *C120F* promoter exhibits a small amount of basal activity. Notably, these data also indicate that background expression is independent of TAEL-N.

As expected, GFP expression increased in response to light in double transgenic embryos (*Tg(C120F:GFP);Tg(ubb:TAEL-N)*) by 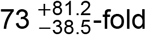 (p=0.0032) (Fig. 6F). In contrast, GFP expression in light-exposed embryos that did not express TAEL-N (*Tg(C120F:GFP)*) increased by only 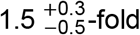. This was a slight but statistically significant difference (p=0.0386). Given that we did not observe any visible GFP fluorescence in illuminated 2 dpf *Tg(C120F:GFP)* embryos (see Fig. 6A), this increase in GFP mRNA levels is unlikely to be functionally relevant.

At 4 dpf, ΔC_T_ values showed elevated background (i.e., dark) GFP expression compared to 2 dpf, presumably due to the ectopic liver expression in this transgenic line (Fig. 6E). The average GFP delta ΔC_T_ values of larvae kept in the dark were 6.2±0.82 for *Tg(C120F:GFP)* and 7.32±0.16 for *Tg(C120F:GFP);Tg(ubb:TAEL-N)* (Fig. 6E). Nevertheless, we could still detect a light-induced increase in GFP expression in *Tg(C120F:GFP);Tg(ubb:TAEL-N)* double transgenic larvae of 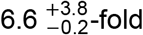 (p=0.0025) (Fig. 6F). Light induction of *Tg(C120F:GFP)* embryos led to an apparent 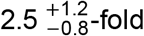 decrease in GFP expression, but this was not statistically significant (p=0.1820).

All together, these data show that in the absence of activated TAEL-N, basal activity of the *C120F* promoter is low and negligibly responsive to light, demonstrating fidelity of the TAEL 2.0 system.

## Discussion

In this study, we describe improvements we have made to a zebrafish-optimized optogenetic expression system called TAEL/C120. In the original TAEL/C120 system, a LOV domain-containing transcription factor (TAEL) is used to drive expression of genes of interest downstream of the C120 regulatory element in response to blue light. The changes we made include adding a C-terminal nuclear localization signal to TAEL (TAEL-N) and coupling C120 regulatory elements with a basal promoter taken from the mouse *Fos* gene (*C120F*). These improvements, collectively referred to as TAEL 2.0, significantly increased both the level and rate of light-induced expression.

Importantly, these improvements allowed us to generate functional stable transgenic lines for TAEL-N expression. Previously under TAEL 1.0, we had difficulties generating such transgenic lines, possibly due to sub-optimal performance of the TAEL transcriptional activator and/or sensitivity of the C120 promoter. We speculate that these deficiencies were overcome in TAEL 1.0 by transiently expressing TAEL by mRNA or plasmid injection, which can deliver many more molecules of TAEL than can be achieved by transgene expression. However, this approach limits the applications for TAEL 1.0 as injections are labor intensive, introduce experimental variability, and often preclude use beyond early embryonic stages. In this study, the improvements we made to both the transcriptional activator (TAEL-N) and promoter (*C120F*) together allowed us to generate functional TAEL-N transgenic lines. Such lines can provide additional spatiotemporal specificity to gene induction, as demonstrated with the *Tg(sox17:TAEL-N)* line (Fig. 4). And, as shown with the *Tg(ubb:TAEL-N)* line (Fig. 5), transgenesis enables usage beyond early embryonic stages, which is not possible with mRNA delivery to the zygote.

Several different basal promoters have been used in zebrafish transgene and enhancer trap constructs, each with different characteristics^15,21,23,24^. For a synthetic expression system such as TAEL, an ideal basal promoter will cause negligible background expression while enabling high rates of transcription following induction. In our original TAEL 1.0 system, the *C120* regulatory element is coupled to a minimal TATA box sequence taken from a mammalian expression vector^6,8^. Although it is easily inducible and causes low background, it is not optimized for use in zebrafish and may thus result in lower than desired expression levels. In this study, we replaced the minimal TATA box with the basal promoter of the mouse *Fos* gene, which was previously used in zebrafish transgenesis^15,21^. This modification alone resulted in more than 40-fold activation following illumination — a 15-fold increase over the original TAEL system (Fig. 1C). Although the *Fos* basal promoter is derived from a gene well-known for its activation in response to neuronal activity^25^, our experiments indicate that coupling this basal promoter to the C120 regulatory sequence imparts several desirable attributes to the TAEL system (fast induction, low background, high amplitude) that extend to the whole organism. For cell type-specific applications, further improvement may be possible by choosing a different basal promoter optimized for that cell type.

TAEL 2.0 joins a growing toolkit for light-controlled gene expression in zebrafish. Other light-gated transcriptional activators shown to function in zebrafish include GAVPO^4,26^, a cryptochrome (CRY2/CIB1)-based system^3^, and a phytochrome (Phy/PIF)-based system^27^. Although all are capable of driving light-induced gene expression, each system possesses distinct qualities that users could leverage for different applications. The phytochrome-based system is responsive to red and far-red light, while TAEL-N, GAVPO, and the cryptochrome-based system are responsive to blue light. The GAVPO, cryptochrome-, and phytochrome-based systems were developed by fusing light-sensitive protein domains to the yeast Gal4 transcriptional activator allowing them to be combined with existing *UAS* transgenic lines. In contrast, EL222, from which TAEL-N is derived, was engineered from an endogenously occurring light-activated bacterial transcription system with its own regulatory element (*C120*), making it orthogonal to Gal4/*UAS* approaches. Both the cryptochrome- and phytochrome-based transcriptional activators operate as heterodimers (CRY2/CIB1 and Phy/PIF, respectively) while TAEL-N functions as a homodimer, potentially simplifying experimental design by having one less component to express. The stability of the activated state of each of these transcriptional activators also varies. Activated GAVPO has a relatively long half-life of approximately 2 hours^4^, making it suitable for “cellular memory” applications while activated EL222 and, presumably, N-TAEL have an estimated half-life of 30 seconds^8^, making it ideal for applications where precise on/off control is desired. In short, the properties of TAEL 2.0 are complementary to these other optogenetic expression systems, and users should feel empowered to choose the system best suited for their intended applications.

With the improvements that we have made, we envision TAEL 2.0 will facilitate a broad range of applications including lineage tracing and precise targeting (spatially and temporally) of gene perturbations. One major advantage of TAEL 2.0 is the extension of these applications beyond early embryonic stages through transgene-directed expression of the TAEL-N transcription factor. This improved zebrafish-optimized light-gated gene expression system should be a broadly useful resource for the zebrafish community.

## Acknowledgements

We thank Anna Reade, Didier Stainier, and Orion Weiner for support and advice during early stages of this work. We thank members of the Woo and Materna labs for providing insightful comments throughout the work and especially on the manuscript. We thank Chris Amemiya for generously providing access to the Lecia stereomicroscope. We thank Roy Hoglund, Emily Slocum, and their staff for excellent zebrafish care. This work was funded by grants from the National Institutes of Health (NIH) (R03 DK106358) and the University of California Cancer Research Coordinating Committee (CRN-20-636896) to S.W. K.S. received support from the NSF-CREST: Center for Cellular and Biomolecular Machines at the University of California, Merced (NSF-HRD-1547848). K.H.G. acknowledges support from NIH (R01 GM106239).

